# miRmedon: confident detection of microRNA editing

**DOI:** 10.1101/774661

**Authors:** Amitai Mordechai, Alal Eran

## Abstract

microRNA (miRNA), key regulators of gene expression, are prime targets for adenosine deaminase acting on RNA (ADAR) enzymes. Although ADAR-mediated A-to-I miRNA editing has been shown to be essential for orchestrating complex processes, including neurodevelopment and cancer progression, only a few human miRNA editing sites have been reported. Several computational approaches have been developed for the detection of miRNA editing in small RNAseq data, all based on the identification of systematic mismatches of ‘G’ at primary adenosine sites in known miRNA sequences. However, these methods have several limitations, including their ability to detect only one editing site per sequence (although editing of multiple sites per miRNA has been reproducibly validated), their focus on uniquely mapping reads (although 20% of human miRNA are transcribed from multiple loci), and their inability to detect editing in miRNA genes harboring genomic variants (although 73% of human miRNA loci include a reported SNP or indel). To overcome these limitations, we developed miRmedon, that leverages large scale human variation data, a combination of local and global alignments, and a comparison of the inferred editing and error distributions, for a confident detection of miRNA editing in small RNAseq data. We demonstrate its improved performance as compared to currently available methods and describe its advantages.

**Availability and implementation:** Python source code is available at https://github.com/Amitai88/miRmedon

## Introduction

Owing to their double stranded structure, miRNA are considered prime targets for A-to-I RNA editing, a site-specific deamination of adenosine to inosine carried out by adenosine deaminase acting on RNA (ADAR) enzymes. miRNA are central regulators of gene expression and their editing has been shown to be essential for orchestrating complex processes such as neuronal development and cancer progression (Behm and Öhman, 2016; Wang and Liang, 2018). A-to-I editing can efficiently alter miRNA biogenesis and/or targeting, thereby leading to global gene expression shifts (Tassinari et al., 2019). Despite their emerging role in human health and disease, very few human miRNA editing sites have been reported (Pinto et al., 2018).

Several computational approaches have been developed for the detection of miRNA editing in small RNAseq data, all based on the identification of systematic mismatches of ‘G’ at primary adenosine sites in known miRNA sequences (Alon et al., 2012; Vesely et al., 2012; Zheng et al., 2016). These methods have several limitations, including the ability to detect only one editing site per sequence (although editing of multiple sites per miRNA has been reproducibly validated), the focus on uniquely mapping reads only (although roughly half of the human miRNA genes are transcribed from multiple loci), and inability to detect editing miRNA harboring genomic variants (although 73% of human miRNA loci include a SNP or indel). To overcome these limitations we developed miRmedon, <u>miR</u>NA <u>m</u>ultiple <u>ed</u>iting detecti<u>on</u>.

A major barrier in confidently detecting editing events in small RNAseq is cross-mapping, namely the likely mapping of a putatively edited miRNA sequence to multiple loci, including those from which it did not originate (de Hoon et al., 2010). For this reason, Alon et al. restricted their analysis to uniquely mapping small RNA reads (Alon et al., 2012). However, this requirement excludes editing of the 20% of all mature human miRNA that could have originated from several genomic loci. A more tolerant approach was taken by Zheng et al. (Zheng et al., 2016) that used a cross-mapping correction method (de Hoon et al., 2010). Yet, this method cannot detect multiple editing sites per miRNA, since considering all possible editing events in mature miRNA aggravates the cross-mapping issue.

To overcome these limitations, we developed miRmedon, a novel framework for the confident detection of multiple editing sites in a single read. miRmedon is based on a combination of local and global alignments to the genome and transcriptome, as well as consideration of large-scale population variation data, and a comparison of the inferred editing distribution with the sequencing error distribution. We demonstrate its improved performance for *de novo* detection of A-to-I miRNA editing in several small RNAseq datasets, as compared to currently available methods.

## Description

### e-miRbase generation

The first step of miRmedon includes the *in-silico* generation of an extended, human variation-aware, edited human miRNA collection. For that purpose, miRbase (Release 21, Griffiths-Jones et al., 2006) was used to simulate all possible editing combinations, in which each miRNA is represented by all possible editing combinations, i.e. with a guanosine in each primary adenosine position. Editing sites mapped to a gnomAD SNP or those within 5bp of a gnomAD indel were excluded (a total of 18.8% of sites, gnomAD release 2.1.1, Karczewski et al., 2019). All simulated edited miRNA sequences were globally aligned to the human genome (GRCh38.p12,) and transcriptome (Gencode 31,) with Seqmap (Jiang and Wong, 2008). Sequences that perfectly matched non-miRNA loci or transcripts were discarded (**Figure 1A**).

**Fig 1.**
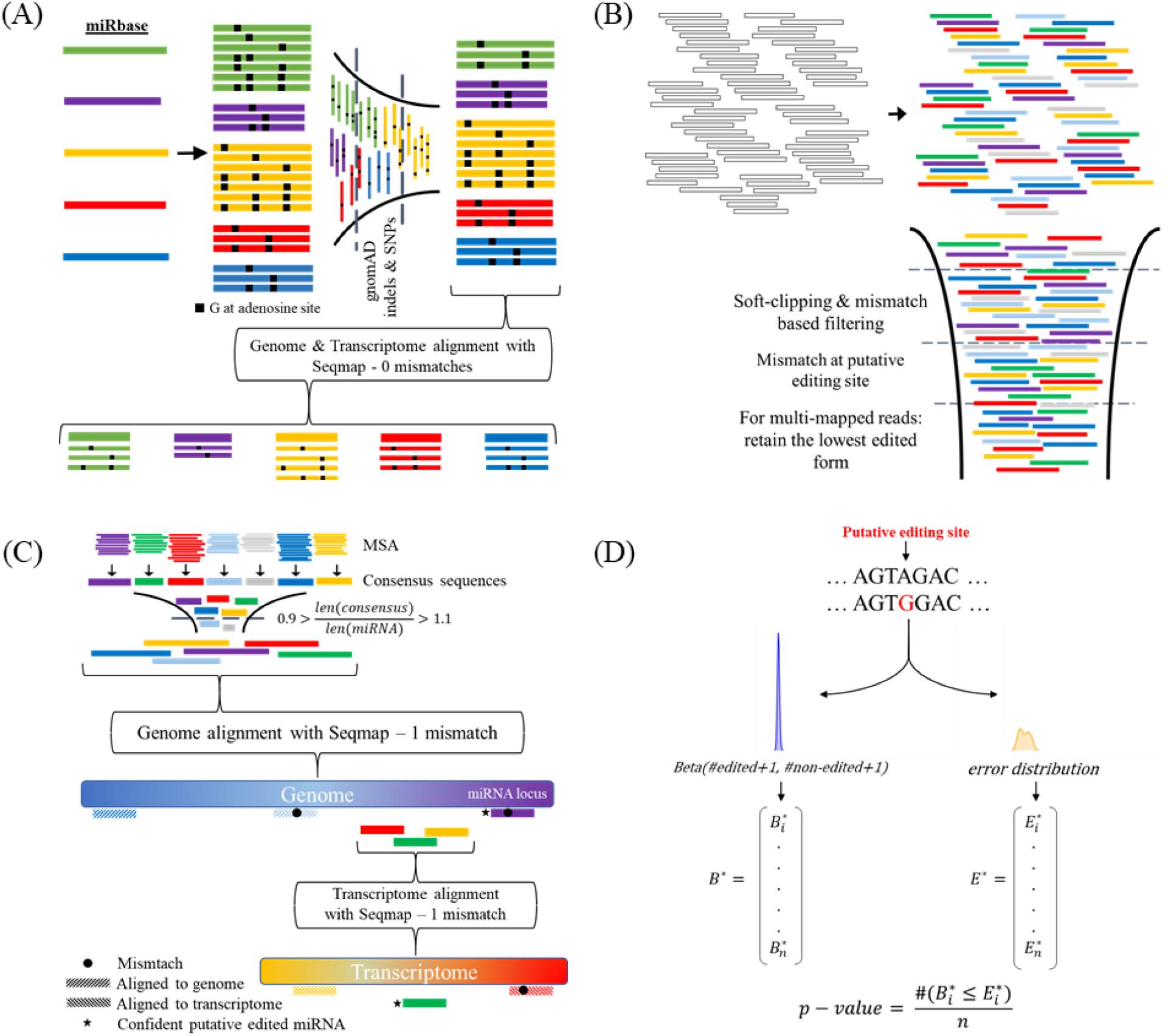
miRmedon pipeline. (**A**) e-miRbase - An extended version of miRbase is generated, in which each mature miRNA sequence is represented by all possible combinations of putative editing. Each edimiRNA sequence was aligned to human genome and transcriptome. Finally, all non-edited miRNAs and speculated edited miRNA sequences were assembled to form e-miRbase. (**B**) Read clustering - Small-RNAseq reads are aligned to e-miRbase in order to identify potentially edited miRNA reads following by several post-alignment filtering steps. (**C**) Read classification –for each reads cluster, a multiple sequence alignment was performed followed by quality-dependent consensus sequence construction. After discarding unappropriate length consensus sequences, the remaining sequences are aligned to reference genome. Completely aligned sequences are disposed, likewise sequences aligned with 1 mismatch to non-miRNA locus. On the contrary, un-aligned sequences and those aligned with 1 mismatch to miRNA locus, are further treated as speculated edited miRNA sequences. Second alignment step is performed aligning all un-aligned sequences to reference transcriptome. Again, all aligned sequences, with up to 1 mismatch, are also discard. (**D**) Inference and quantification – for each putative editing site, a beta distribution with the parameters α=1+number of reads with a ‘G’ at site, and β=1+number of ‘A’ at putative site + 1. Likewise, a second distribution is formed by calculating error sequencing probability extracted from ‘G’ base calls at the putative editing site. A bootstrapped distribution with *n* samples was formed for the beta distribution (*B*\*), as well as for sequencing error distribution (*E*\*), and a p-value was calculated as the number of times 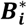 is less than or equal to 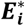 divided by *n*.

### Read clustering

Trimmed and quality inspected small RNAseq reads were locally aligned to e-miRbase using STAR (version 2.6.1a) (Dobin et al., 2013). Further post-alignment filtering steps are taken. First, reads that were soft-clipped more than X bp (adjustable parameter) at the ends are removed. In a similar manner, the number of soft-clipped bases + number of mismatches could be restricted. Second, reads aligning with a non-A/G mismatch to the putative editing site are removed. Finally, multi-mapped reads that aligned to non-edited and edited reads are considered non-edited. Also, reads that mapped to several edited form, are counted as the lowest edited form. At that stage and after read quantification, lowly expressed clusters of both edited and non-edited miRNAs are ruled-out, while each reference sequence for a multi-mapped read is counted as *1/number of loci*. (**Figure 1B**)

### Read classification

Consensus sequences are generated for each reads cluster correspond to edited miRNA incorporating both sequence and Phred quality score (**supplementary materials and methods**). Next, consensus sequences of shorter or longer length than mature miRNA reference sequence length (0.9 < ratio < 1.1) are filtered. Remaining consensus sequences are aligned to human genome (GRCh38.p12 (https://www.ncbi.nlm.nih.gov/grc)), with 1 mismatch allowed, using Seqmap (Jiang and Wong, 2008). Edited miRNA forms whose representative sequences found aligned to locations not recognized as miRNA loci are discarded. Sequences which not found aligned to human genome are aligned to human transcriptome (Gencode version 31 (Harrow et al., 2012)) with the same procedure. Again, clusters representative sequences found aligned to human transcriptome are not further considered as putative edited miRNA sequences. (**Figure 1C**)

### Editing site inference and editing level quantification

For each speculated editing sites, a posterior editing density is a beta distribution with parameters α = number of reads with G mapped to the site + 1, and β = number of reads with A mapped to the site + 1 is formed. Additionally, error probabilities, as computed from Phred quality scores at each putative editing site, were collected to form a second distribution. Monte-Carlo simulation was performed in order to evaluate the confidence level for each putative editing site. Specifically, samples are randomly drawn from the two distributions and compared. P value, namely the probability of observing guanosine in each primary adenosine position due to sequencing error, is calculated as the number of incidences in which error distribution drawn sample was equal to or greater than editing rate distribution drawn sample, divided by the number of resamples. (**Figure 1D**)

## Advantages over existing methods

### Improved sensitivity

**a**. By generating all possible combinations of substituting ‘A’ to ‘G’ at non variance-prone adenosine sites, miRmedon allows the discovery of multiple *prima facie* editing events in a sole read. As a result, miRmedon enables the detection of linked editing sites, i.e. editing events with high probability of been observed simultaneously in single read. Moreover, previous proposed methods do not consider reads containing additional non-reference mismatches arising by possible sequencing errors or SNPs (besides A/G mismatches) as edited miRNA, thus fail to estimate the real editing rate. In certain cases, substantial existent signal of A-to-I editing events will deemed as not-significant, i.e. as a sequencing error artifact, due to this bias.

**b.** RNA modifications are overrepresented at the 3’ end of mature miRNA (Burroughs et al., 2010), thus impeding the alignment process when using global aligner like Bowtie (Langmead et al., 2009), which have been utilized in previous suggested pipelines. In addition, small RNA seq data frequently demonstrating low-quality calls at read’s ends, further contributing to this predicament. Recognizing that, the mapping process (of both edited and non-edited miRNA reads) improve by employing STAR’s local alignment mode to align small-RNA seq reads to e-miRbase, which allows for soft-clipping at read’s ends. This utility spares the suggested transversal trimming of bases at the reads 3’ end (Alon et al., 2012).

**c.** Our proposed statistical inference for discriminant substantial A-to-I editing signal from sequencing error empower detection ability effectively to discover even lowly-edited sites, thereby propitious also for small cohorts’ studies.

### Improved specificity

**a.** Since genomic SNV could incorrectly considered as A-to-I RNA editing event, thus all current tools discard SNPs recorded at dbSNP (Sherry et al., 2001). Our pipeline leverages large scale population variation data (gnomAD) to reduce the risk for false-positive results in pursuance of SNPs. Furthermore, we exclusively exclude also all primary adenosine sites located at ±5 of indels indicated by gnomAD.

**b**. The cross-mapping issue (de Hoon et al., 2010), which become more grievous by considering all possible combinations of putative editing sites, is addressed by aligning consensus sequences of each presumed edited miRNA reads cluster to the genome with Seqmap (Jiang and Wong, 2008), an aligner designed for finding all possible loci where a read could originate from. In this manner, and by generating quality-dependent and reference-free consensus sequences, the most likely representative sequence of each cluster is obtained – hence, enabling accurate mapping and identification of speculated edited miRNAs as distinct genomic entities. For this purpose, The GRCh38.p12 genome assembly of all regions was used, including reference chromosomes, scaffolds, assembly patches and haplotypes. Moreover, consensus sequences are also aligned to the transcriptome, in order to neglect any possibility of mRNA fragment been considered as edited miRNA.

## Results

To demonstrate the enhanced performance of miRmedon we exploited pooled human brain small RNAseq data analyzed by Alon et al. (Alon et al., 2012) used them for introducing what became predominantly the standard tool for miRNA editing detection. While Alon et al. discovered only 18 significant editing sites analyzing that dataset, miRmedon revealed 52 significant editing sites (**Supplementary Table 1**). 12 out of 52 sites are overlapping with the founding of Alon et al. (**Supplementary Fig S1**), while 5 sites of the remaining weren’t considered due to low supported reads and higher threshold set in our analysis. Additional editing site detected by Alon et al. was an editing site at the 6^th^ position of hsa-miR-376b-3p, nevertheless our pipeline detected an editing site at the 6^th^ position of hsa-miR-376a-3p. Both miRNAs differ only at the last 3 bases, thus several reads could potentially map to both miRNAs. Worth to mention that Alon at al. didn’t report the 6^th^ position of hsa-miR-376a-3p as an editing site.

As it was argued, miRmedon enables the detection of multiple editing sites in a single read. Indeed, several such reads were identified, for example the 12^th^ and 17^th^ position of hsa-let-7f-5p (**Supplementary Fig S2**). While the 17^th^ position found edited independently of the 12th position, disregarding reads which contain G at both editing sites will results with an extremely low editing rate of hsa-let-7f-5p 17^th^ position (∼0.14%) – which be probably inferred as a sequencing error by Alon at al. pipeline. Nevertheless, miRmedon inferred both 12^th^ and 17^th^ as editing sites with editing levels of 1.06% and 1.21% respectively.

We compared our results to a comprehensive survey conducted by Pinto et al. (Pinto et al., 2018) employing Alon et al. pipeline to analyze 10,593 samples of several tissues and conditions resulting with a list of 58 reliable editing sites within 55 mature miRNAs. In addition, Pinto’s survey including a list of 129 editing sites found in previous studies within 98 mature miRNAs. Essentially, those 129 sites could be collapsed into 125 editing sites, considering identical mature miRNAs which potentially originated from different hairpins. Likewise, the 58 editing sites indicated by Pinto et al. are 57 mature miRNA sites. Considering mature miRNA expression of the examined sample, there are 75 editing sites potential for detection, while 39 of them noted by Pinto et al. as highly confident sites. We found an overlap of 19 sites with 75 potentially previous reported editing sites, while 16 were included in Pinto’s high confident editing sites (Supplementary Fig S2). Conclusively, miRmedon revealed 33 novel editing sites. It should be noted that the ability of our tool to detect Pinto’s confident sites which were not identified by Alon et al. isn’t dependent on editing level (**Supplementary Fig S3**).

Examining the flanking nucleotides of the 52 editing sites detected in pooled human brain sample for the 5’-UAG-3’ preferred motif of ADAR enzymes (Kleinberger and Eisenberg, 2010) we observed certain bias for ‘U’ at the −1 position of the editing site (17 sites), nevertheless G was much more represented at the +1 position (23 sites). Additionally, a depletion of G at the −1 position (5 sites) is observed, as expected from editing site (Kleinberger and Eisenberg, 2010). Even by inspecting only the 36 sites which weren’t indicated by Pinto et al. as high confident editing sites, a strong signal for ADAR preference motif is still maintained with 12 U and 18 G observed in −1 and +1 position respectively, together with a depletion of G at the −1 position (4 sites). (**Supplementary Fig S4**).

Next, we have taken Alon et al. strategy to endeavor validating editing sites detected in pooled human brain tissue, by examine the effect of overexpressing ADAR1 or ADAR2 in glioblastoma cell-line and compare editing rates in each putative editing site to control cells. First, we analyzed U87 glioblastoma cell-line transiently over-expressing ADAR1. While 29 out of 52 putative edited miRNAs were expressed in control cells, only one editing site showed slightly elevated editing level in treated cells (5.7% Vs. 3.7%). Second, we followed the same procedure to analyze U118 cell-line stably over-expressing ADAR2. While 27 putative edited miRNA which found in pooled brain sample were expressed in U118 cell-line, 7 found valid as ADAR2 substrates with no significant editing found in control cells. Remarkably, 3 sites which found valid by this analysis are novel not yet reported A-to-I editing sites. (**Supplementary Fig S5**).

## Conclusion

miRmedon detects human A-to-I miRNA editing with better sensitivity and specificity as compared to currently available approaches. Thus, employing it for *de-novo* detection of miRNA editing in small RNAseq data can be harnessed to identify novel editing sites and gain much-needed insights into the implications of miRNA editing in human health and disease.

## References

Alon, S., Mor, E., Vigneault, F., Church, G.M., Locatelli, F., Galeano, F., Gallo, A., Shomron, N., Eisenberg, E., 2012. Systematic identification of edited microRNAs in the human brain. Genome Res. 22, 1533–1540. https://doi.org/10.1101/gr.131573.111

Behm, M., Öhman, M., 2016. RNA Editing: A Contributor to Neuronal Dynamics in the Mammalian Brain. Trends Genet. 32, 165–175. https://doi.org/10.1016/j.tig.2015.12.005

Burroughs, A.M., Ando, Y., de Hoon, M.J.L., Tomaru, Y., Nishibu, T., Ukekawa, R., Funakoshi, T., Kurokawa, T., Suzuki, H., Hayashizaki, Y., Daub, C.O., 2010. A comprehensive survey of 3’ animal miRNA modification events and a possible role for 3’ adenylation in modulating miRNA targeting effectiveness. Genome Res. 20, 1398–1410. https://doi.org/10.1101/gr.106054.110

de Hoon, M.J.L., Taft, R.J., Hashimoto, T., Kanamori-Katayama, M., Kawaji, H., Kawano, M., Kishima, M., Lassmann, T., Faulkner, G.J., Mattick, J.S., Daub, C.O., Carninci, P., Kawai, J., Suzuki, H., Hayashizaki, Y., 2010. Cross-mapping and the identification of editing sites in mature microRNAs in high-throughput sequencing libraries. Genome Res. 20, 257–264. https://doi.org/10.1101/gr.095273.109

Dobin, A., Davis, C.A., Schlesinger, F., Drenkow, J., Zaleski, C., Jha, S., Batut, P., Chaisson, M., Gingeras, T.R., 2013. STAR: ultrafast universal RNA-seq aligner. Bioinforma. Oxf. Engl. 29, 15–21. https://doi.org/10.1093/bioinformatics/bts635

Griffiths-Jones, S., Grocock, R.J., van Dongen, S., Bateman, A., Enright, A.J., 2006. miRBase: microRNA sequences, targets and gene nomenclature. Nucleic Acids Res. 34, D140–D144. https://doi.org/10.1093/nar/gkj112

Harrow, J., Frankish, A., Gonzalez, J.M., Tapanari, E., Diekhans, M., Kokocinski, F., Aken, B.L., Barrell, D., Zadissa, A., Searle, S., Barnes, I., Bignell, A., Boychenko, V., Hunt, T., Kay, M., Mukherjee, G., Rajan, J., Despacio-Reyes, G., Saunders, G., Steward, C., Harte, R., Lin, M., Howald, C., Tanzer, A., Derrien, T., Chrast, J., Walters, N., Balasubramanian, S., Pei, B., Tress, M., Rodriguez, J.M., Ezkurdia, I., van Baren, J., Brent, M., Haussler, D., Kellis, M., Valencia, A., Reymond, A., Gerstein, M., Guigó, R., Hubbard, T.J., 2012. GENCODE: The reference human genome annotation for The ENCODE Project. Genome Res. 22, 1760–1774. https://doi.org/10.1101/gr.135350.111

Jiang, H., Wong, W.H., 2008. SeqMap: mapping massive amount of oligonucleotides to the genome. Bioinforma. Oxf. Engl. 24, 2395–2396. https://doi.org/10.1093/bioinformatics/btn429

Karczewski, K.J., Francioli, L.C., Tiao, G., Cummings, B.B., Alföldi, J., Wang, Q., Collins, R.L., Laricchia, K.M., Ganna, A., Birnbaum, D.P., Gauthier, L.D., Brand, H., Solomonson, M., Watts, N.A., Rhodes, D., Singer-Berk, M., Seaby, E.G., Kosmicki, J.A., Walters, R.K., Tashman, K., Farjoun, Y., Banks, E., Poterba, T., Wang, A., Seed, C., Whiffin, N., Chong, J.X., Samocha, K.E., Pierce-Hoffman, E., Zappala, Z., O’Donnell-Luria, A.H., Minikel, E.V., Weisburd, B., Lek, M., Ware, J.S., Vittal, C., Armean, I.M., Bergelson, L., Cibulskis, K., Connolly, K.M., Covarrubias, M., Donnelly, S., Ferriera, S., Gabriel, S., Gentry, J., Gupta, N., Jeandet, T., Kaplan, D., Llanwarne, C., Munshi, R., Novod, S., Petrillo, N., Roazen, D., Ruano-Rubio, V., Saltzman, A., Schleicher, M., Soto, J., Tibbetts, K., Tolonen, C., Wade, G., Talkowski, M.E., Consortium, T.G.A.D., Neale, B.M., Daly, M.J., MacArthur, D.G., 2019. Variation across 141,456 human exomes and genomes reveals the spectrum of loss-of-function intolerance across human protein-coding genes. bioRxiv 531210. https://doi.org/10.1101/531210

Kleinberger, Y., Eisenberg, E., 2010. Large-scale analysis of structural, sequence and thermodynamic characteristics of A-to-I RNA editing sites in human Alu repeats. BMC Genomics 11, 453. https://doi.org/10.1186/1471-2164-11-453

Langmead, B., Trapnell, C., Pop, M., Salzberg, S.L., 2009. Ultrafast and memory-efficient alignment of short DNA sequences to the human genome. Genome Biol. 10, R25. https://doi.org/10.1186/gb-2009-10-3-r25

Pinto, Y., Buchumenski, I., Levanon, E.Y., Eisenberg, E., 2018. Human cancer tissues exhibit reduced A-to-I editing of miRNAs coupled with elevated editing of their targets. Nucleic Acids Res. 46, 71–82. https://doi.org/10.1093/nar/gkx1176

Sherry, S.T., Ward, M.-H., Kholodov, M., Baker, J., Phan, L., Smigielski, E.M., Sirotkin, K., 2001. dbSNP: the NCBI database of genetic variation. Nucleic Acids Res. 29, 308–311.

Tassinari, V., Cesarini, V., Silvestris, D.A., Gallo, A., 2019. The adaptive potential of RNA editing-mediated miRNA-retargeting in cancer. Biochim. Biophys. Acta BBA - Gene Regul. Mech., mRNA modifications in gene expression control 1862, 291–300. https://doi.org/10.1016/j.bbagrm.2018.12.007

Vesely, C., Tauber, S., Sedlazeck, F.J., von Haeseler, A., Jantsch, M.F., 2012. Adenosine deaminases that act on RNA induce reproducible changes in abundance and sequence of embryonic miRNAs. Genome Res. 22, 1468–1476. https://doi.org/10.1101/gr.133025.111

Wang, Y., Liang, H., 2018. When microRNAs meet RNA editing in cancer: A nucleotide change can make a difference. BioEssays News Rev. Mol. Cell. Dev. Biol. 40. https://doi.org/10.1002/bies.201700188

Zheng, Y., Ji, B., Song, R., Wang, S., Li, Ting, Zhang, X., Chen, K., Li, Tianqing, Li, J., 2016. Accurate detection for a wide range of mutation and editing sites of microRNAs from small RNA high-throughput sequencing profiles. Nucleic Acids Res. 44, e123. https://doi.org/10.1093/nar/gkw471

