## Supplementary file for "miRmedon: confident detection of microRNA editing"

### Supplementary materials and methods

#### Reference free quality dependent consensus sequence

First, multiple sequence alignment is applied for each read cluster $R$ of speculated edited miRNA form using Mafft (Katoh et al., 2002; Katoh and Standley, 2013) utilizing the full length read. For each aligned read $A_i \in A$ in each position $j$, a discrete probability function representing the probability to observe each of the calls in the set $N = \{A, T, C, G, N/-\}$ is formed, with '-' represents a gap generated in the MSA process. Provided that $A_{i,j}$ is an un-ambiguous nucleotide (A, T, C or G), the probability $p$ for the observed base is calculated from the Phred score, while the probability to observe each of the alternative nucleotides is $p/3$. The probability to observe an ambiguous character (N/-) is set to $\varepsilon$. On the contrary, if $A_{i,j}$ results as an 'N' or '-', the probability for an ambiguous character is set to $1 - \varepsilon$, and the probability to observe each of the legitimate nucleotides is $(1 - \varepsilon)/4$. The probability to observe the character $N_k \in N$ is given by equation1 (Genest and Zidek, 1986) which used to calculate the merged probability function for to observe $N_k$ in position $j$ based on all reads. Finally, the character in each position $j$ of the consensus sequence is determine as the most likely base $N_k$ as calculated by the merged probability function. In cases which there are several equally likely characters, the consensus sequence in that certain position would be 'N'.

**(eq. 1)** $$\frac{\prod_i P(A_{i,j} = N_k)}{\prod_i P(A_{i,j} = N_k) + \prod_i P(A_{i,j} \neq N_k)}$$

#### Small-RNA seq data

Publicly available small-RNA seq data was downloaded from NCBI Sequence Read Archive (SRA). List of SRR entries are listed in below table:

| SRR | Description |
|---|---|
| SRR095854 | Small-RNA sequencing of pooled human brain sample |
| SRR1615247 | Stable transfection of inactive ADAR2 to U118 cell-line followed by small-RNA sequencing |
| SRR1615246 | Stable transfection of ADAR2 to U118 cell-line followed by small-RNA sequencing |
| SRR346127 | Small-RNA sequencing of U87 cells |
| SRR346128 | Transient transfection of ADAR1 to U87 followed by small-RNA sequencing |

#### Quality control

Small RNA reads were first subject to quality control using FastQC (Andrews S, 2010). Unless already trimmed, adapters were removed using cutadapt (Martin, 2011) following by quality-based filtering with Trimmomatic (Bolger et al., 2014).

#### Supplementary results

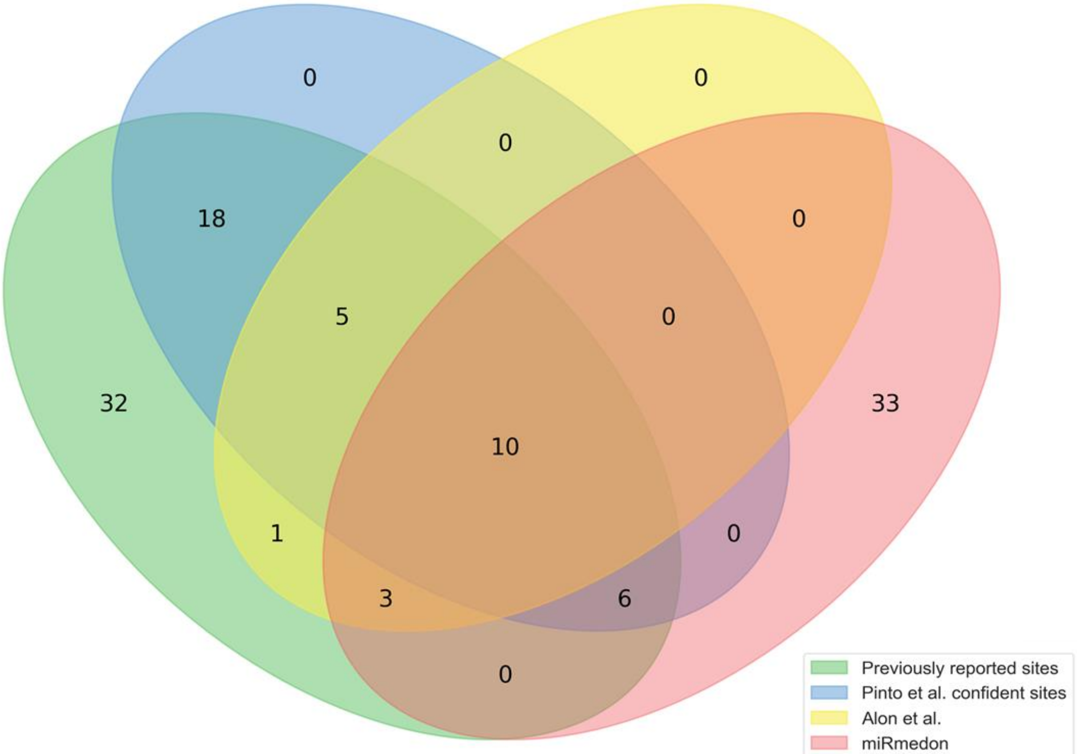


**Fig S2.** Intersection between *de-novo* detected sites in pooled human brain sample (pink) with sites detected by Alon et al. pipeline (Alon et al., 2012) (yelow) using the same dataset, previously reported editing sites (green) and a list of 58 reliable editing sites as indictated by Pinto et al (Pinto et al., 2018) (blue).

| Read sequence | Count |
| --- | --- |
| TGAGGTAGTAGGTTGTGTAGTTT | 6 |
| TGAGGTAGTAGGTTGTGTAGTT | 45 |
| TGAGGTAGTAGGTTGTGTAGT | 22 |
| TGAGGTAGTAGGTTGTGTAT | 4 |
| TGAGGTAGTAGGTTGTGAAG | 1 |
| TGAGGTAGTAGGTTGTGTAC | 1 |
| TGAGGTAGTAGGTTGTGTAGTTA | 6 |
| TGAGGTAGTAGGTTGTGTA | 33 |
| TGAGGTAGTAGGTTGTGTAA | 68 |
| TGAGGTAGTAGGTTGTGTAGTA | 1 |
| AGGTAGTAGGTTGTGTAA | 1 |
| TGAGGTAGTAGGTTGTGTAG | 9 |
| TGAGGTAGTAGGTTGTGTAGA | 2 |
| TGAGGTAGTAGGTTGTGTAGTTG | 1 |
| TGAGGTAGTAGATTGTGTAGTTA | 1 |
| TGAGGTAGTAGATTGTGTAGTTT | 2 |
| TGAGGTAGTAGATTGTGTAGT | 3 |
| TGAGGTAGTAGTTTGTGTA | 1 |
| TGAGGTAGTAGATTGTGTAGTT | 17 |
| TGAGGTAGTAGATTGTGTAG | 1 |
| TGAGGTAGTAGATTGTGTAGTA | 2 |

**Fig S3**. Reads mapped to edited form of hsa-let-7f-5p. Both detected editing events (marked in red), at position 12 and 17, are novel A-to-I editing sites. While editing site at the 17$^{th}$ position was detected separately, no editing was obseved at the 12$^{th}$ position by itself. Presumably, previous suggested pipelines wouldn't detect even the 17$^{th}$ position as an editing sites, due to very low editing rate (~0.14%) unless counting editing events occuring in conjugation with an editing at the 12$^{th}$ nucelotide.

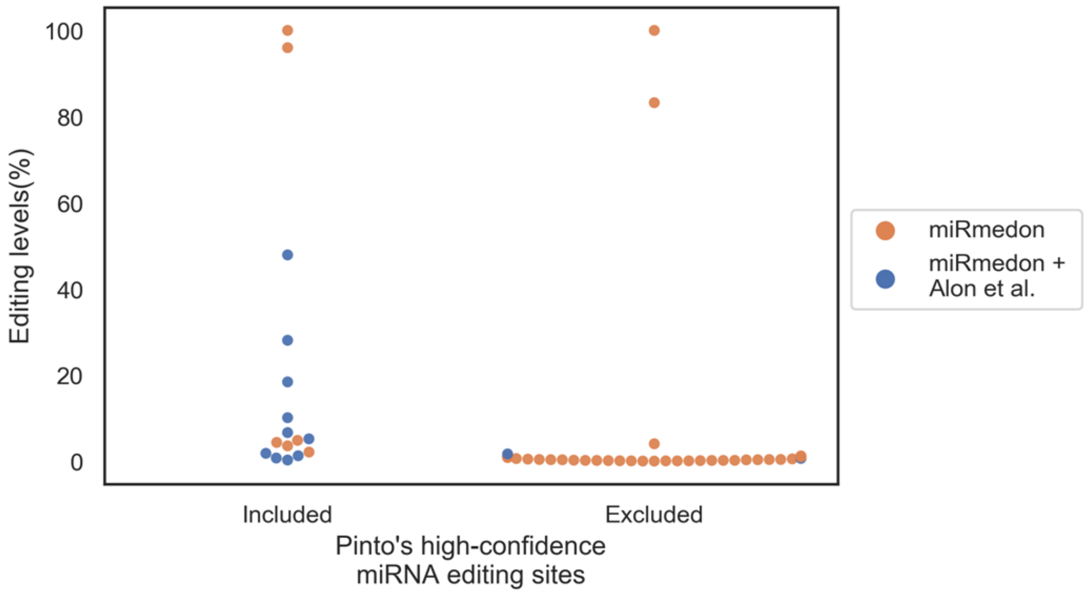


**Fig S4.** Editing rate of 52 editing sites detected by miRmedon in pooled human brain sample classified to sites included in Pinto et al. high confident sites and those excluded. Featured here the ability of our tool, comparing to Alon et al. pipeline (Alon et al., 2012), in the detection reliable editing sites in small scale datasets independently of editing rate.

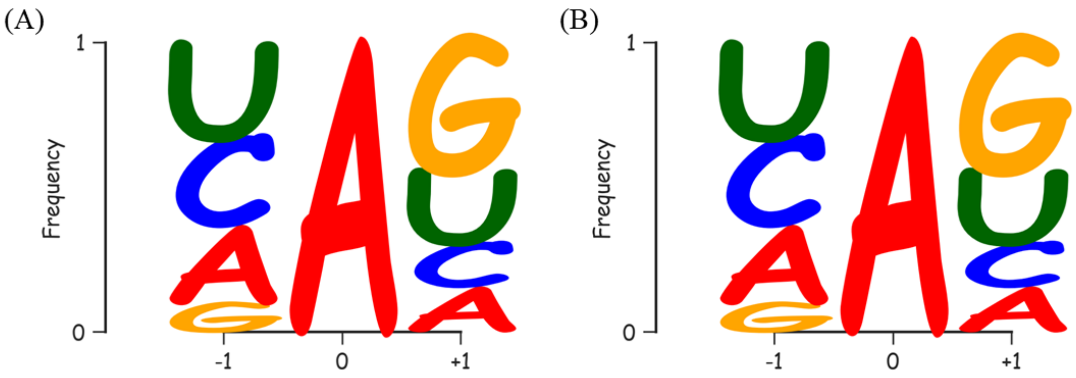


**Fig S5.** Motif preference of putative editing sites detected in pooled human brain sample. Both motifs, of all 52 sites (**A**) and 36 sites which weren't included in Pinto et al. 58 reliable sites (**B**), exhibit the expected up-stream and down-stream nucleotides propensity of genuine editing sites with a certain bias to U and depletion of G at -1 position and a bias to G at the +1 position of the editing site.

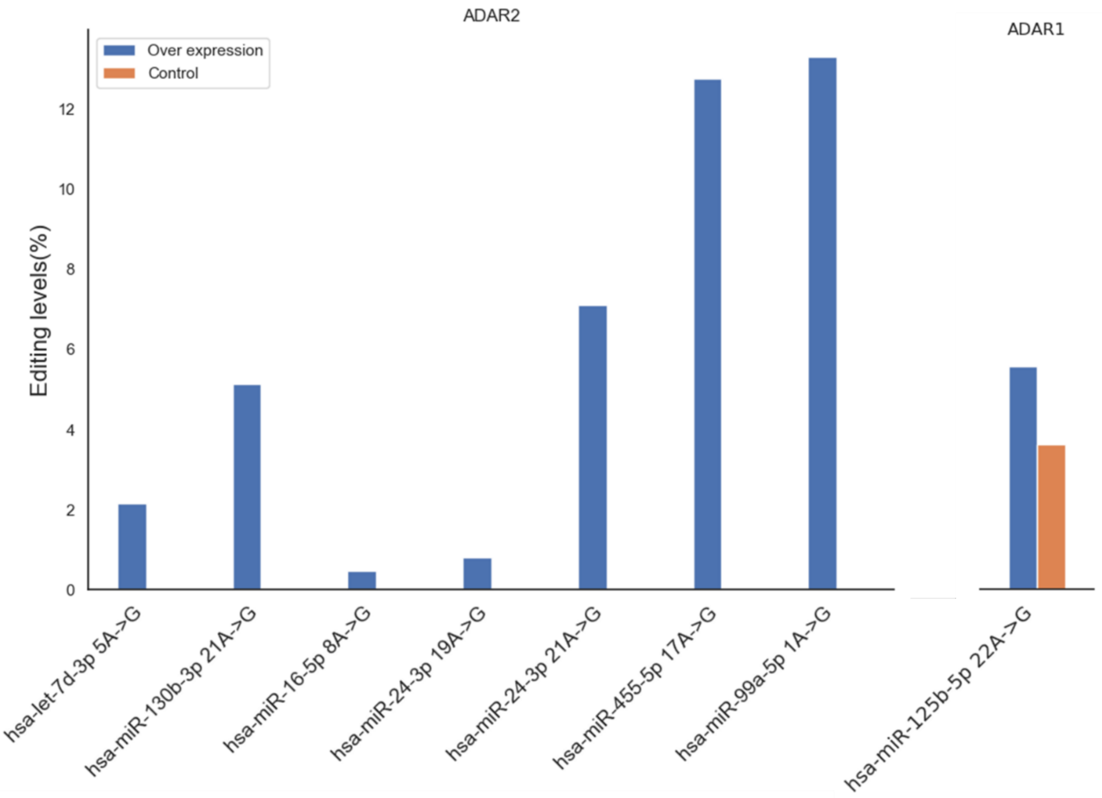

**Fig S6**. Editing levels of *de-novo* detected editing sites validated by in-vitro overexpression of ADAR1 and ADAR2 in globlastoma cell-lines (U87 and U118 respectively). Blue bars represent editing levels of ADAR overexpressing cells, while orange bars represent editing levels of control cells. while U87 cells overexpressing ADAR1 have shown some elevated editing level compared to control cells at the 22th position of hsa-miR-125-5p (5.7% Vs 3.7%), all sites validated as ADAR2 targets weren't exhibit any significant signal for G at the putatative editing sites.

### References


Alon, S., Mor, E., Vigneault, F., Church, G.M., Locatelli, F., Galeano, F., Gallo, A., Shomron, N., Eisenberg, E., 2012. Systematic identification of edited microRNAs in the human brain. Genome Res. 22, 1533–1540. https://doi.org/10.1101/gr.131573.111

Andrews S, 2010. FastQC: a quality control tool for high throughput sequence data. Available online at: http://www.bioinformatics.babraham.ac.uk/projects/fastqc.

Bolger, A.M., Lohse, M., Usadel, B., 2014. Trimmomatic: a flexible trimmer for Illumina sequence data. Bioinformatics 30, 2114–2120. https://doi.org/10.1093/bioinformatics/btu170

Dobin, A., Davis, C.A., Schlesinger, F., Drenkow, J., Zaleski, C., Jha, S., Batut, P., Chaisson, M., Gingeras, T.R., 2013. STAR: ultrafast universal RNA-seq aligner. Bioinforma. Oxf. Engl. 29, 15–21. https://doi.org/10.1093/bioinformatics/bts635

Genest, C., Zidek, J.V., 1986. Combining Probability Distributions: A Critique and an Annotated Bibliography. Stat. Sci. 1, 114–135. https://doi.org/10.1214/ss/1177013825

Jiang, H., Wong, W.H., 2008. SeqMap: mapping massive amount of oligonucleotides to the genome. Bioinforma. Oxf. Engl. 24, 2395–2396. https://doi.org/10.1093/bioinformatics/btn429

Katoh, K., Misawa, K., Kuma, K., Miyata, T., 2002. MAFFT: a novel method for rapid multiple sequence alignment based on fast Fourier transform. Nucleic Acids Res. 30, 3059–3066.

Katoh, K., Standley, D.M., 2013. MAFFT Multiple Sequence Alignment Software Version 7: Improvements in Performance and Usability. Mol. Biol. Evol. 30, 772–780. https://doi.org/10.1093/molbev/mst010

Langmead, B., Trapnell, C., Pop, M., Salzberg, S.L., 2009. Ultrafast and memory-efficient alignment of short DNA sequences to the human genome. Genome Biol. 10, R25. https://doi.org/10.1186/gb-2009-10-3-r25

Martin, M., 2011. Cutadapt removes adapter sequences from high-throughput sequencing reads. EMBnet.journal 17, 10–12. https://doi.org/10.14806/ej.17.1.200

Pinto, Y., Buchumenski, I., Levanon, E.Y., Eisenberg, E., 2018. Human cancer tissues exhibit reduced A-to-I editing of miRNAs coupled with elevated editing of their targets. Nucleic Acids Res. 46, 71–82. https://doi.org/10.1093/nar/gkx1176
